# Association of Ddx5/p68 protein with the upstream erythroid enhancer element (EHS1) of the Klf1 gene

**DOI:** 10.1101/743435

**Authors:** Xiaoyong Chen, Sanjana Pillay, Felix Lohmann, James J Bieker

**Affiliations:** Department of Cell, Developmental, and Regenerative Biology, Mount Sinai School of Medicine, New York, NY; Black Familly Stem Cell Institute, Mount Sinai School of Medicine, New York, NY; Tisch Cancer Institute, Mount Sinai School of Medicine, New York, NY; Mindich Child Health and Development Institute, Mount Sinai School of Medicine, New York, NY

**Keywords:** transcription factor, biochemical affinity isolation, erythropoiesis, gene expression, transcription

## Abstract

EKLF/KLF1 is an essential transcription factor that plays a global role in erythroid transcriptional activation. It’s own regulation is of interest, as it displays a highly restricted expression pattern, limited to erythroid cells and its progenitors. Here we use biochemical affinity purification to identify the Ddx5/p68 protein as an activator of KLF1 by virtue of its interaction with the erythroid-specific DNAse hypersensitive site upstream enhancer element (EHS1). We postulate that its range of interactions with other proteins known to interact with this element render it part of the enhanseosome complex critical for optimal expression of KLF1. These individual interactions provide quantitative contributions that, in sum, establish high level activity of the KLF1 promoter and suggest they can be selectively manipulated for clinical benefit.

## Introduction

The mechanisms by which intracellular transcriptional regulators interact to direct hematopoietic stem cells towards a particular lineage and exert control in target expression remains a major question in cellular and molecular studies (1-4). Analysis of the erythroid lineage has led to the successful characterization of regulators that act as transcription factors and establish the proper network to generate erythroid-specific expression. However, in many cases it remains unresolved how these factors are themselves regulated.

One way to address these issues is to isolate genomic clones that code for the gene-specific regulators and determine their own cis-acting regulatory elements and the trans-acting factors that bind to them. Based on this notion, we have been studying the regulation of erythroid Krüppel-like factor (EKLF;KLF1 (5)), an erythroid-enriched transcription factor that is intimately involved in global regulation of downstream erythroid-specific gene expression by binding to cognate CCMCRCCCN elements (6-9). A number of functional properties and expression characteristics of KLF1 make it of interest to study its regulation.

First, KLF1 expression remains tissue-restricted throughout early development and in the adult. Its onset is strictly limited to the mesodermal, primitive erythroid cells that populate the blood islands of the extraembryonic yolk sac at the early headfold stage (E7.5), switching by E9.5 to definitive cells within the hepatic primordia, and then to the red pulp of the adult spleen (10). It is expressed in the erythroblastic island macrophage, an in vivo niche that supplies a supportive environment for developing red cells (11-13). Early in hematopoietic differentiation KLF1 is expressed at low levels in the MPP and it retains an expression pattern restricted to the CMP and MEP as its transcript levels increase prior to eventual segregation to erythroid progeny (14-16).

Second, KLF1’s activation target repertoire has expanded beyond the ß-like globin gene locus to include protein-stabilizing, heme biosynthetic pathway, red cell membrane protein, cell cycle, and transcription factor genes in both primitive and definitive erythroid cells (6-8,17). Relatedly, links have been established between mutant or haploinsufficient levels of KLF1 and altered human hematology and anemia (6,18,19), as some genes are uniquely sensitive to haploinsufficiency (20-22).

We have shown that a 950 base pair region, located just upstream of the KLF1 transcription initiation site, is sufficient to generate erythroid-specific expression in transient assays (23). This region exhibits the most significant homology upon a seven species alignment of 30 kB of surrounding genomic DNA (24), and harbors erythroid-restricted DNAse hypersensitive sites. One of these sites (erythroid hypersensitive site 1; EHS1) behaves as a very strong enhancer, which in conjunction with the proximal promoter (25-27) accounts for KLF1’s tissue-specific expression (23). The importance of this short region has been verified *in vivo* (28-30). We have suggested a two-tiered mechanism for transcriptional regulation of KLF1, with GATA2 and SMAD5 proteins initially generating low transcript levels, followed by high quantities of KLF1 transcript after GATA1 protein is produced (24). Coupled to inclusion of the DEK protein into this complex along with erythroid histone marks (31) and the RIOK2 protein (32), EHS1 can now be considered a well-characterized erythroid enhancer.

Our identification of DEK relied on a biochemical affinity purification to a region of EHS1 denoted as ‘oligo2’ region of EHS1 (31). In the present study, we have identified a second protein that also binds to that region in vitro and contributes to optimal enhancement activity by EHS1 in vivo.

## Results

We had utilized a magnetic bead approach based on DNA affinity technology, coupled with mass spec analysis of the selected, associated proteins, to isolate proteins that bind the “oligo2” sequence within EHS1. This enabled identification of the “35kD” protein as DEK (31). We next focused our attention on the oligo2-enriched protein at ∼60kD. MALDI/TOF isolation of peptides and their sequence analysis yielded a particularly significant hit with the p68/Ddx5 protein (Figure 1).

**Figure 1.**
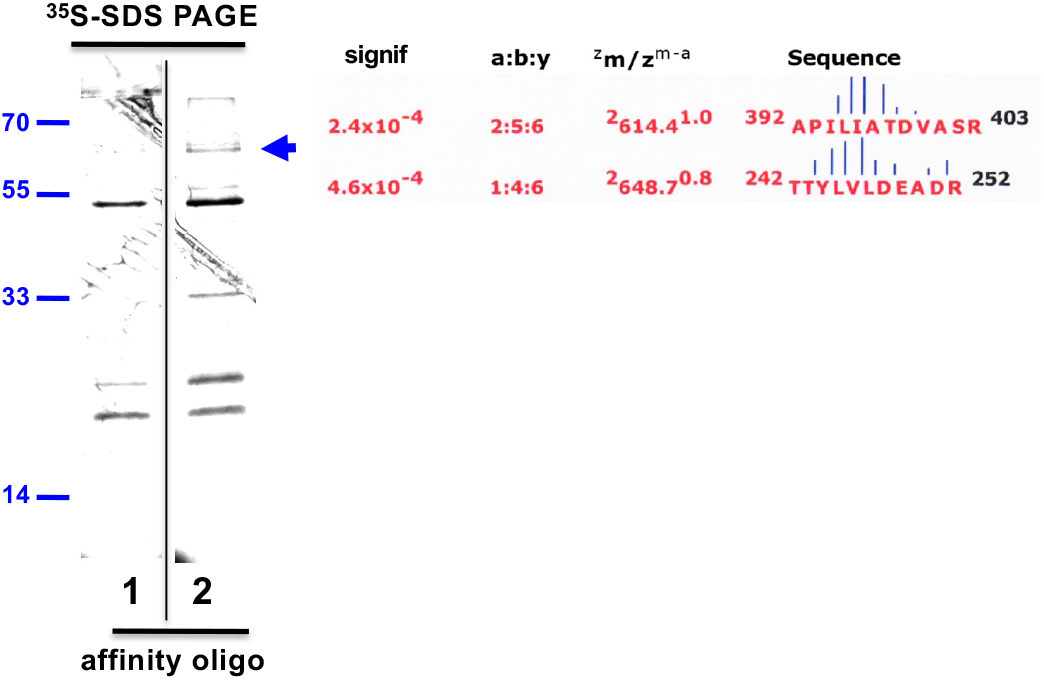
Analysis of affinity-purified proteins. DNA binding proteins that interact with oligo2 were monitored by SDS/PAGE analysis of ^35^S methionine-labeled proteins after oligo1 (negative control (23)) or oligo2 affinity purification. Two non-adjacent lanes from the same gel are shown; a vertical line marks the boundary between the two. Purified material derived from ∼5×10E6 cell equivalents are shown. Arrowhead on right marks the protein unique to oligo 2 preparation that is the focus of this study, along with the mass spec data (Sonar MS/MS analysis: a:b:y ratio is of the fragmentation ions; ^z^m/z^m–a^ is the ^charge^mass/charge^measured minus calculated^ mass; the vertical bar between amino acid pairs indicates the ion intensity within the peptide fragment (59)) and peptide sequences used in its identification; lines on left indicate molecular weight as indicated (kD).

There is a considerable literature on Ddx5 protein and its interactions with nucleic acids (33,34).

Ddx5 is a generally expressed ‘DEAD box’ family protein that interacts with DNA via chromatin, and plays a role in a range of RNA processing complexes. Ddx5 contains an ATP-dependent RNA helicase activity that is an important part of the Drosha complex (35,36), although evidence suggests that this activity is dispensable for its transcriptional regulatory function (33,34). Disruption leads to mouse embryonic lethality by E11.5 (35).

Ddx5 is well expressed within erythroid cells at all stages of differentiation, and in progenitors (Figure 2A). One interesting aspect in this case (compared to DEK) is that there is no evidence that p68/Ddx5 interacts with a specific DNA sequence. Regardless, we performed a chromatin immunoprecipitation (ChIP) test for Ddx5 interaction with the EHS1 region in MEL cells, and find that it binds to a high level (relative to IgG). Use of a variant MEL line that has deleted the endogenous ‘oligo2’ site shows that Ddx5 binding decreases ∼2.5 fold (Figure 2B). These data are consistent with prior published studies on the importance of the ‘oligo2’ site, which showed that it is required for optimal binding of transcriptional activators such as P300 and Tal1, and that its removal leads to a quantitative (2-fold) reduction in production of KLF1 RNA (31).

**Figure 2.**
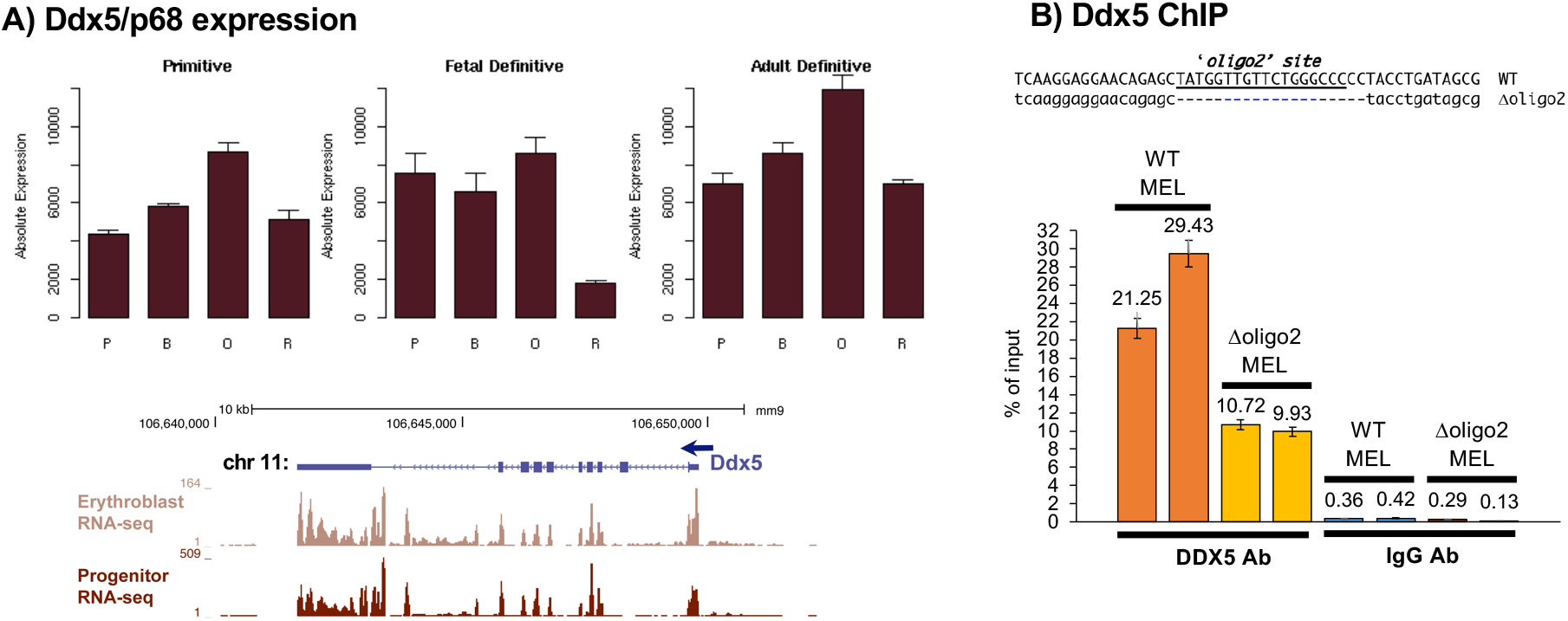
Erythroid expression of Ddx5/p68. **A)** *Top*, data from the Erythron database (63) shows expression levels of Ddx5/p68 during mouse primitive and definitive (fetal liver or bone marrow) differentiation. P=proerythroblasts, B=basophilic erythroblasts, O=orthochromatiphilic erythroblasts, R=reticulocytes. *Bottom*, browser data of RNA-seq expression in erythroid progenitors and erythroblast cells. **B)** In vivo chromatin immunoprecipitation of Ddx5 protein or isotype IgG binding to the EHS1 site in WT MEL cells and MEL cells containing a genomic deletion centered on oligo2 (oligo2 sequence (23) is underlined, and the extent of deletion on Δoligo2 MEL line (31) is as indicated). Results from two experiments each performed in triplicate are shown.

Of relevance to this point, Ddx5 interacts with P300 and CBP (37), and is part of a hematopoietic cell complex that forms with Scl/Tal1 (38), a protein already known to interact with the KLF1 EHS1 region (39,40). Ddx5 also interacts with Smad5 (36,41), a protein we and others have shown is critical for KLF1 induction (24,42). As a result, it is relatively straightforward to envision the Ddx5 protein’s role in KLF1 transcriptional activation via EHS1 (Figure 3).

**Figure 3.**
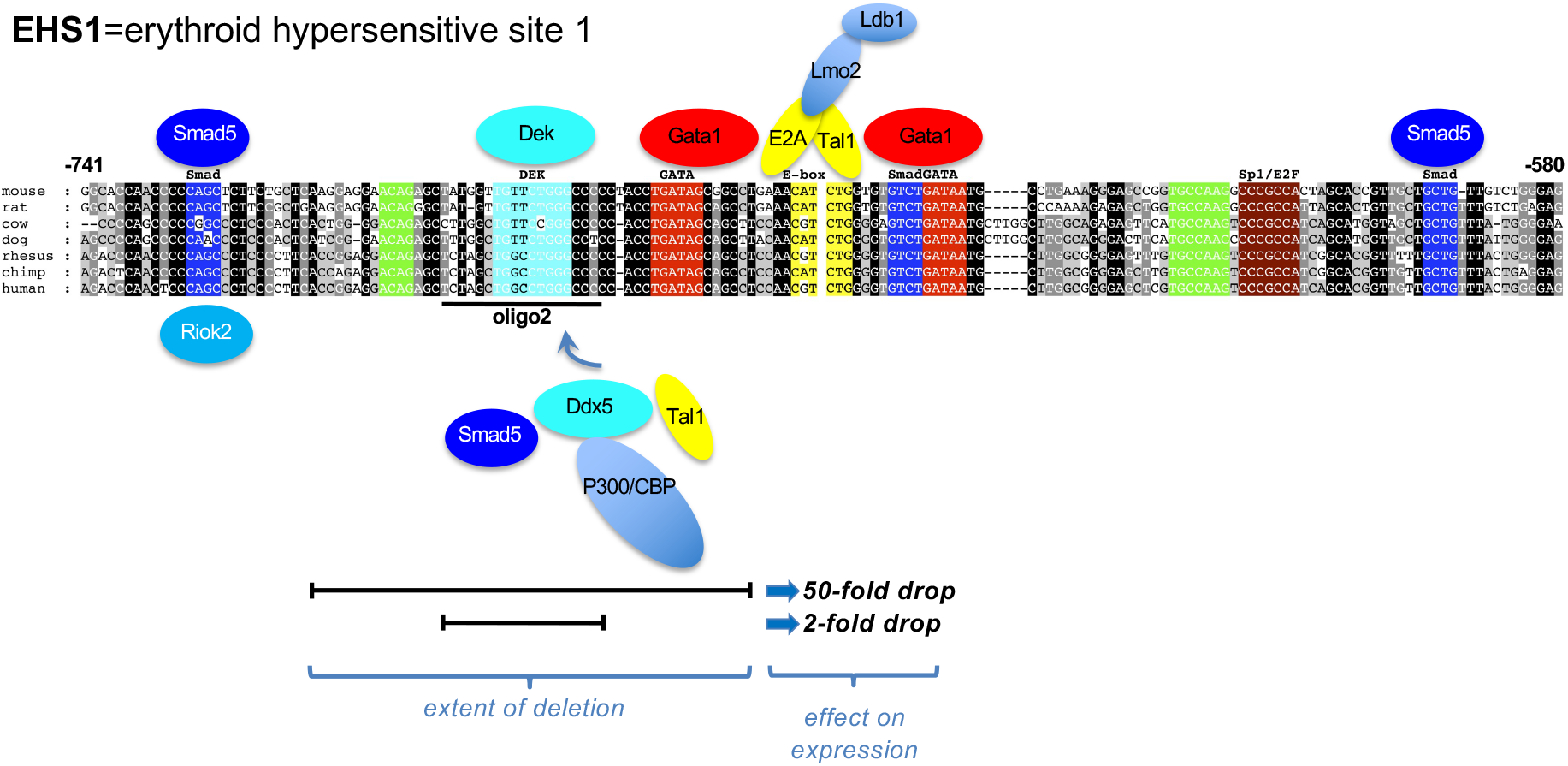
Sequence homology of KLF1 EHS1. Detailed layout of conserved *Klf1* cis-regulatory elements at the upstream enhancer (EHS1) (24,31). Blocks of conserved sequence homology between seven mammalian species are color-coded for their transcription factor binding sites, highlighting Smad, DEK, Gata-Ebox-Gata sites; location of oligo2 used in this study (based on (23)) is as indicated. Also drawn in are schematics of potential enhanceosome components at EHS1: *Above* includes the known multicomponent Gata/Tal1/Lmo2/Ldb1 multi-protein complex (based on (3,39,64-66)); *below* brings in known Ddx5 protein interactions as discussed in the text, including the recently-described interaction of Riok2 with this region (32). Also indicated are the extent of deletions that lead to altered levels of Klf1 expression: the 49 bp deletion of -715 to -666 (23) that leads to a 50-fold drop (30), and the 18 bp deletion of oligo2 (23) that leads to a 2-fold drop (31).

## Discussion

We have identified p68/Ddx5 as a novel protein that interacts with the ‘oligo2’ region of EHS1. Interestingly, Ddx5 performs much of its function via the long noncoding (lnc) RNA molecule SRA (43,44), an interaction that is important for insulator control of gene expression (45). Ddx5 plays a necessary role in skeletal muscle differentiation, in part by complexing with and stabilizing the activating myoD/Brg1 interaction and aiding the recruitment of TBP and RNA pol II to muscle gene promoters (46). SRA has been shown to regulate global erythroid gene expression in primary human cells and cell lines (47).

Based on these properties, we hypothesize that Ddx5, in a complex with other proteins at EHS1 that likely form an enhanseosome (Figure 3), plays a directive role in establishing a 3-dimensional structure at the KLF1 genomic locus. This is consistent with the chromatin architecture and long-range KLF1 genetic region interaction points observed in erythroid cells (48). Related to this point, Ddx5 also interacts with CTCF (45), a protein that plays a critical role in higher order chromatin structure and enhancer insulation (49-51). RNA pull-down of SRA recovers Ddx5 and CTCF in a complex (47,52). CTCF is already a well-known player in erythroid gene regulation, as it is required for proper looping at the ß-globin, SCL/TAL1, and myb transcription units (53-57).

A critical aspect from the extended studies of KLF1 gene regulation is that although removal of the complete EHS1 site leads to a dramatic 50-fold reduction in KLF1 expression, removal of the 18-bp ‘oligo2’ site leads to a more nuanced effect (Figure 3). These properties demonstrate it is possible to identify genetic modifications that quantitatively alter KLF1 levels, thus mimicking haploinsufficiency and potentially providing a clinically useful approach to relieve hemoglobinopathies by its effect of altering ß-like globin gene regulation (20-22).

### Experimental Procedures

32DEpo1 and MEL cells were grown as previously described (23,58). The MEL line containing a genetic deletion of the oligo2 sequence has been described (“indel 8”; (31)).

EHS1-binding proteins were purified as previously described (31), focused on ‘oligo2’ (23). Briefly, M280 magnetic Dynabeads coupled with streptavidin (Life Technologies) were incubated with biotinylated oligo2 to create the oligo-Dynabeads. The 32DEpo1 cell lysate (23) from >10 liters of cells was added and incubation continued at room temperature with rotation. Using a magnetic apparatus, the beads were washed to remove most of the non-specific binding proteins before elution in high salt. Gel shift assays were used throughout the procedure to estimate enrichment (31). Eluted protein was dialyzed and concentrated by acetone precipitation prior to SDS-PAGE electrophoresis. Proteins were visualized by staining with colloidal blue and bands of interest were excised. Mass spec analyses of the gel slices were performed by the Rockefeller University Protein and DNA Technology Center using the Sonar MS/MS search engine coupled with statistical scoring methods (59).

Chromatin immunoprecipitation (ChIP) was performed using the anti-Ddx5 antibody (Bethyl A300-523A; (60)). 10 × 10^6^ cells (per replicate) were crosslinked in 1% formaldehyde at room temperature for 10 min and quenched with 0.125M glycine for 5 mins at room temperature followed by 15 mins on ice. The cell pellet was washed twice with PBS to remove excess formaldehyde and resuspended in 1mL of MNase lysis buffer (0.2% NP-40, 10mM NaCl & 10mM Tris pH 8.0) and kept on ice for 15 mins. Cells were spun and resuspended in 500uL of MNase lysis buffer with 0.1M CaCl_2_ and 2uL of MNase (20U/uL) and incubated for 10 min at 37°C. 10uL of 0.1M EDTA was added to stop the reaction and the cells were spun down (4°C) and resuspended in 400uL of RIPA buffer (Cell Signaling Tech) with 1X protease inhibitor cocktail (Roche) and sonicated in a Bioruptor device (22 pulses of 40s on and 60s off at 21 amplitude). The cells were diluted with 1X RIPA buffer to a total volume of 1000uL and centrifuged at 13000 rpm, and 50uL of the was saved as input (20%). The rest of the supernatant was incubated with the anti-Ddx5 antibody (5ug per replicate) and rabbit IgG (Life Technologies) overnight at 4°C on a rotating stand. Protein A Dynabeads (Thermo Scientific) were added the following day and incubated for another 2hrs at 4°C on a rotating stand. The beads were collected using a magnetic stand and the supernatant was discarded. The beads were resuspended and first washed with 1X RIPA buffer (3 times, 5 min each) and then once in TSEII (0.1% SDS, 10% Triton X-100, 0.5M EDTA, 1M Tris pH 8 and 5M NaCl) and twice in TSEIII buffer (1% NP-40, 1% Na-deoxycholate, 0.5M EDTA, 1M Tris pH 8 and 0.25M LiCl). Final washes were done in TE buffer (0.5M EDTA and 1M Tris pH 8) and the DNA was then eluted from the beads at 65°C for 20 min in 200uL of elution buffer (0.2M NaCl, 1% SDS and 1X TE). After the incubation the beads were separated using a magnet and the eluate was transferred to a fresh tube and crosslinked using proteinase K overnight at 65°C. Next day, DNA was purified by phenol:chloroform extraction and the resulting pellet was resuspended in 30uL (IP) and 50uL (input). For qPCR, both the ChIP and input was diluted 1:3, and 3uL of this was used for amplification per reaction (10uL total). The primer pair for quantitation was designed using IDT software and is as follows: 5’GCTCAGACCTCAACACAACA (forward) and 5’TGTCTGATGATGCCTGAAAGG (reverse).

Databases used as part of this study include transcription factor (61,62), and erythroid expression (63) sources.

## Data availability

all data are contained within the manuscript.

## Acknowledgements

This work was supported by NIH grants DK048721 and DK046865 to JJB.

